# Range reduction of the Oblong Rocksnail, *Leptoxis compacta*, shapes riverscape genetic patterns

**DOI:** 10.1101/2020.05.12.090662

**Authors:** Aaliyah D. Wright, Nicole L. Garrison, Ashantye’ S. Williams, Paul D. Johnson, Nathan V. Whelan

**Author notes:** Corresponding Authors: Aaliyah D. Wright^1^, Tuskegee University, Tuskegee, Alabama, USA, Email address, Nathan V. Whelan^2,3^, 203 Swingle Hall, Auburn, AL 36849, USA.

## Abstract

Many freshwater gastropod species face extinction, including 79% of species in the family Pleuroceridae. The Oblong Rocksnail, *Leptoxis compacta*, is a narrow range endemic pleurocerid from the Cahaba River basin in central Alabama that has seen rapid range contraction in the last 100 years. Such a decline is expected to negatively affect genetic diversity in the species. However, precise patterns of genetic variation and gene flow across the restricted range of *L. compacta* are unknown. This lack of information limits our understanding of human impacts on the Cahaba River system and Pleuroceridae. Here, we show that *L. compacta* has likely seen a species-wide decline in genetic diversity, but remaining populations have relatively high genetic diversity. We also report a contemporary range extension compared to the last published survey. *Leptoxis compacta* does not display an isolation by distance pattern, contrasting patterns seen in many riverine taxa. Our findings also indicate that historical range contraction has resulted in the absence of common genetic patterns seen in many riverine taxa like isolation by distance as the small distribution of *L. compacta* allows for relatively unrestricted gene flow across its remaining range despite limited dispersal abilities. Two collection sites had higher genetic diversity than others, and broodstock sites for future captive propagation and reintroduction efforts should utilize sites identified here as having the highest genetic diversity. Broadly, our results support the hypothesis that range contraction will result in the reduction of species-wide genetic diversity, and common riverscape genetic patterns cannot be assumed to be present in species facing extinction risk.

## Introduction

Freshwater gastropods of the United States suffer one of the highest imperilment rates of any taxonomic group in North America (Johnson et al., 2013). Despite being critical components of many freshwater ecosystems, freshwater gastropods are grossly understudied compared to freshwater fish, mussels, and crayfish (Covich, Palmer & Crowl, 1999; Huryn, Benke & Ward, 1995; Strong et al., 2008). This creates a situation where desperately needed conservation efforts are hindered by a lack of information (Johnson et al., 2013). For example, data on the current range of many freshwater gastropods is lacking (Lydeard et al., 2004), but conservation assessments and effective management plans require detailed historical and contemporary range data (Potter & Thomas, 1983; USFWS, 2018). Population genetics data on freshwater gastropods are also needed to inform management efforts and provide basic understanding of freshwater ecosystems (Lysne et al., 2008).

The freshwater gastropod family Pleuroceridae is one group that suffers from a high imperilment rate (79%) and little research attention (Brown, Lang & Perez, 2008; Johnson et al., 2013; Perez & Minton, 2008). Pleurocerids are found east of the Rocky Mountains in North America, with most of their diversity concentrated in the southeastern United States (Lydeard & Mayden, 1995; Strong & Köhler, 2009). Pleurocerids lack a highly vagile veliger larval stage seen in many aquatic gastropod groups, and they are thought to move large distances only when washed downstream (Whelan et al., 2019; Whelan, Johnson & Harris, 2015). Only one study has been published on landscape and conservation genetics of pleurocerids, and that study focused exclusively on a single species, *Leptoxis ampla* (Whelan et al., 2019). Many freshwater species, including *L. ampla*, display common riverscape genetic patterns such as increased genetic diversity in downstream populations and isolation by distance (Hughes, Schmidt & Finn, 2009; Paz-Vinas et al., 2015). However, few studies have tested for such patterns in riverine species that have undergone drastic range reduction, and no such study has been done for a range restricted pleurocerid.

One pleurocerid in need of more research is the Oblong Rocksnail, *Leptoxis compacta* (Figs. 1,2). This species is a narrow range endemic known historically from the middle Cahaba River and a single tributary in central Alabama, USA (Fig. 2; Goodrich, 1922). Until recently, *Leptoxis compacta* was considered extinct as it had not been collected, or at least identified correctly, from 1935 to 2011 (Goodrich, 1941; Johnson et al., 2006; Whelan, Johnson & Harris, 2012). As early as 1941, the decline of *L. compacta* was documented (Goodrich, 1941), and the species now occupies less than 5% of its historical range (Fig. 2; Whelan, Johnson & Harris, 2012). As a narrow range endemic with few historical collections, little is known about the species aside from recent survey efforts and limited life history data (Whelan, Johnson & Harris, 2012). Yet, the rediscovery of *L. compacta* in 2011 resulted in an emergency petition to list the species under the US Endangered Species Act (Kurth, 2017). For management agencies to assess the status of *L. compacta* and design effective conservation plans, detailed survey work and population genetics research are required. Modern population genomic tools such as restriction site associated DNA sequencing (RAD-seq) can provide data that will enhance *L. compacta* management options (Andrews et al., 2016). As a result of having a narrow range along a single river path, an effective recovery strategy for *L. compacta* will likely require reintroduction efforts to previously occupied habitat(s). Maintaining genetic diversity of imperiled species is important for mitigating extinction risk (Frankham, 2005; Frankham, 2010), and reintroduction efforts will require detailed population genetics data to inform broodstock selection for maximizing genetic diversity of captively reared offspring.

**Figure 1:**
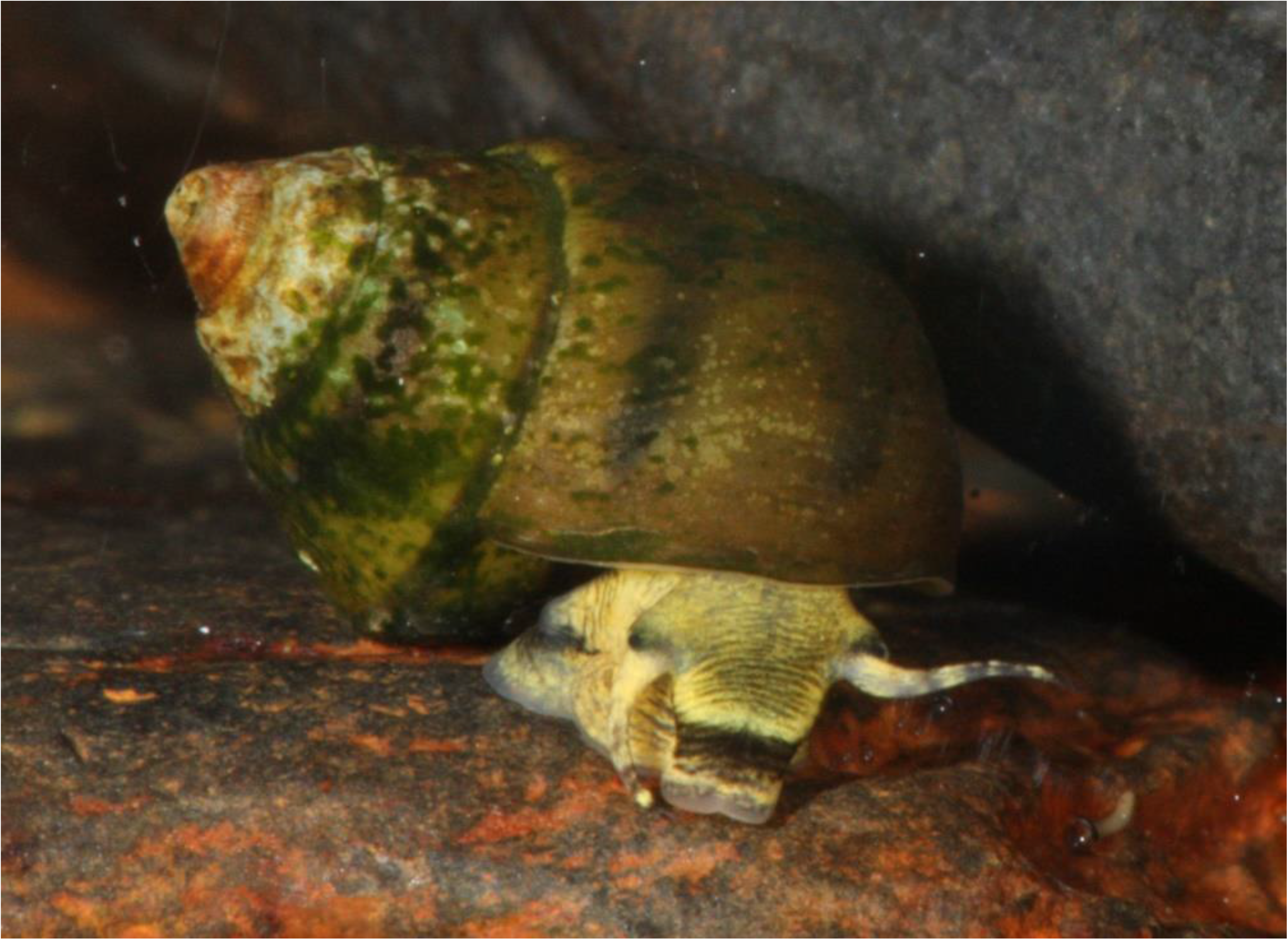
Photograph of live *L. compacta*. Photo Credit: Thomas Tarpley, ADCNR.

**Figure 2:**
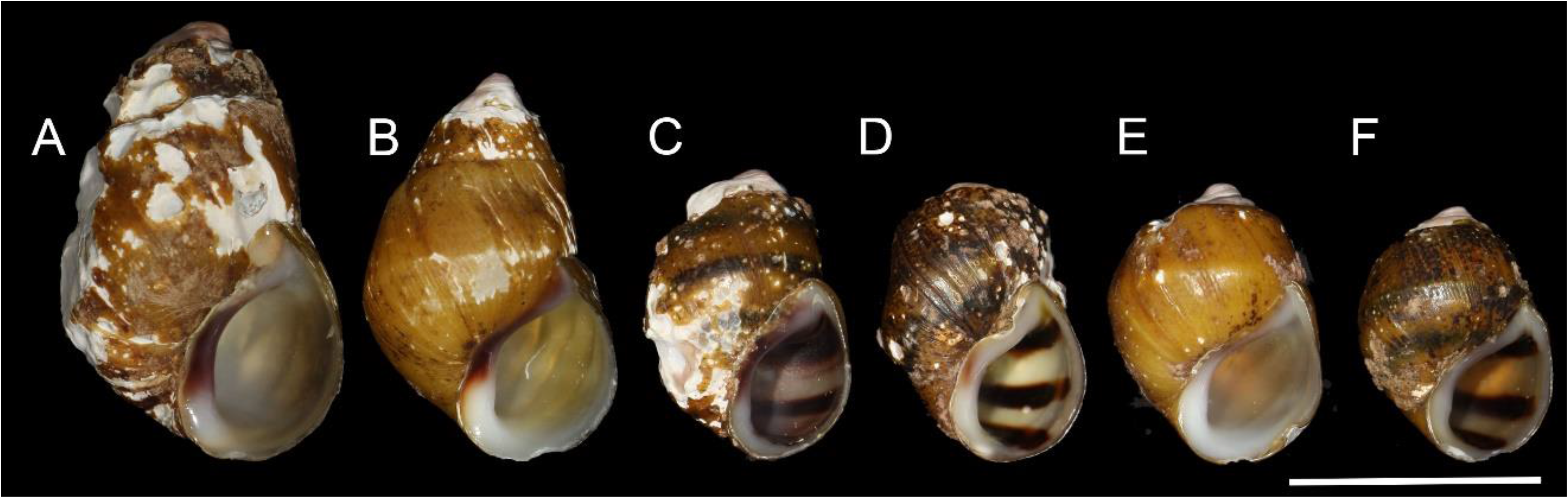
Shells of representative individuals that we sequenced. A) Cahaba River at Canoe Launch, B) Cahaba River at Booth’s Ford, C) Cahaba River above Shades Creek, D-F) Cahaba River at old Marvel slab. Scale bar = 1 cm

In this study, we generated a dataset of thousands of single nucleotide polymorphisms (SNPs) to answer questions about conservation and riverscape genetics of *L. compacta*. Given the drastic range decline suffered by *L. compacta*, we set out to test the following hypotheses: 1) *Leptoxis compacta* has undergone a severe genetic bottleneck and 2) genetic diversity of *L. compacta* is considerably lower than *L. ampla*, a sympatric and wider ranging species. We also examined how genetic diversity of *L. compacta* varies across its current range, specifically assessing whether broad patterns seen in many other riverine taxa like isolation by distance and strong genetic structure are seen in *L. compacta*.

## Materials & Methods

### Sample Collection

*Leptoxis compacta* was collected during two trips to the Cahaba River in June 2018 and June 2019. We performed survey work at four sites, and all sites except Cahaba River above Shades Creek were outside the previously documented contemporary range of *L. compacta* (Fig. 3; Whelan et al. 2012). At each location, individuals were collected by hand and identified in the field. Despite being a narrow range endemic that has undergone distributional decline, *L. compacta* was locally abundant where found. Based on qualitative observations, we sampled less than 1% of the population, making our sampling negligible to species survival. Twenty specimens from each site were transported live to the lab, sacrificed following Fukuda et al., (2008), and placed in 96-100% ethanol until tissue clips could be taken. Specimens were collected under an Alabama Department of Conservation and Natural Resources Educational Scientific Collections Permit (License #2019100990068680) or as an agent of the state (P.D. Johnson). All shells have been cataloged separately to correspond to associated molecular data and deposited at the Auburn University Museum of Natural History (AUMNH 45652-45690; Table 1, Supplementary Table 1).

**Table 1:** Summary statistics and AUMNH catalog numbers of *L. compacta* at each collection site.

| Collection Site | Private<br>alleles | H<sub>o</sub> (sd) | H<sub>e</sub> (sd) | A<sub>r</sub> (sd) | Π (sd) | F<sub>is</sub> | AUMNH # |
| --- | --- | --- | --- | --- | --- | --- | --- |
| Cahaba River at Booth's Ford | 43 | 0.1568 (0.1349) | 0.1801 (0.1222) | 1.8241 (0.2742) | 0.1855 (0.1261) | 0.1319 | 45691-45709 |
| Cahaba River at canoe launch | 32 | 0.1046 (0.1552) | 0.1045 (0.1421) | 1.4511 (0.4518) | 0.1075 (0.1459) | 0.0134 | 45709-45729 |
| Cahaba River above Shades Creek | 262 | 0.1363 (0.1334) | 0.1779 (0.1245) | 1.8072 (0.2883) | 0.1829 (0.1281) | 0.1934 | 45671-45690 |
| Cahaba River at Old Marvel Slab | 28 | 0.0963 (0.1400) | 0.0981 (0.1343) | 1.4606 (0.4387) | 0.1010 (0.1382) | 0.0226 | 45652-45670 |

**Figure 3:**
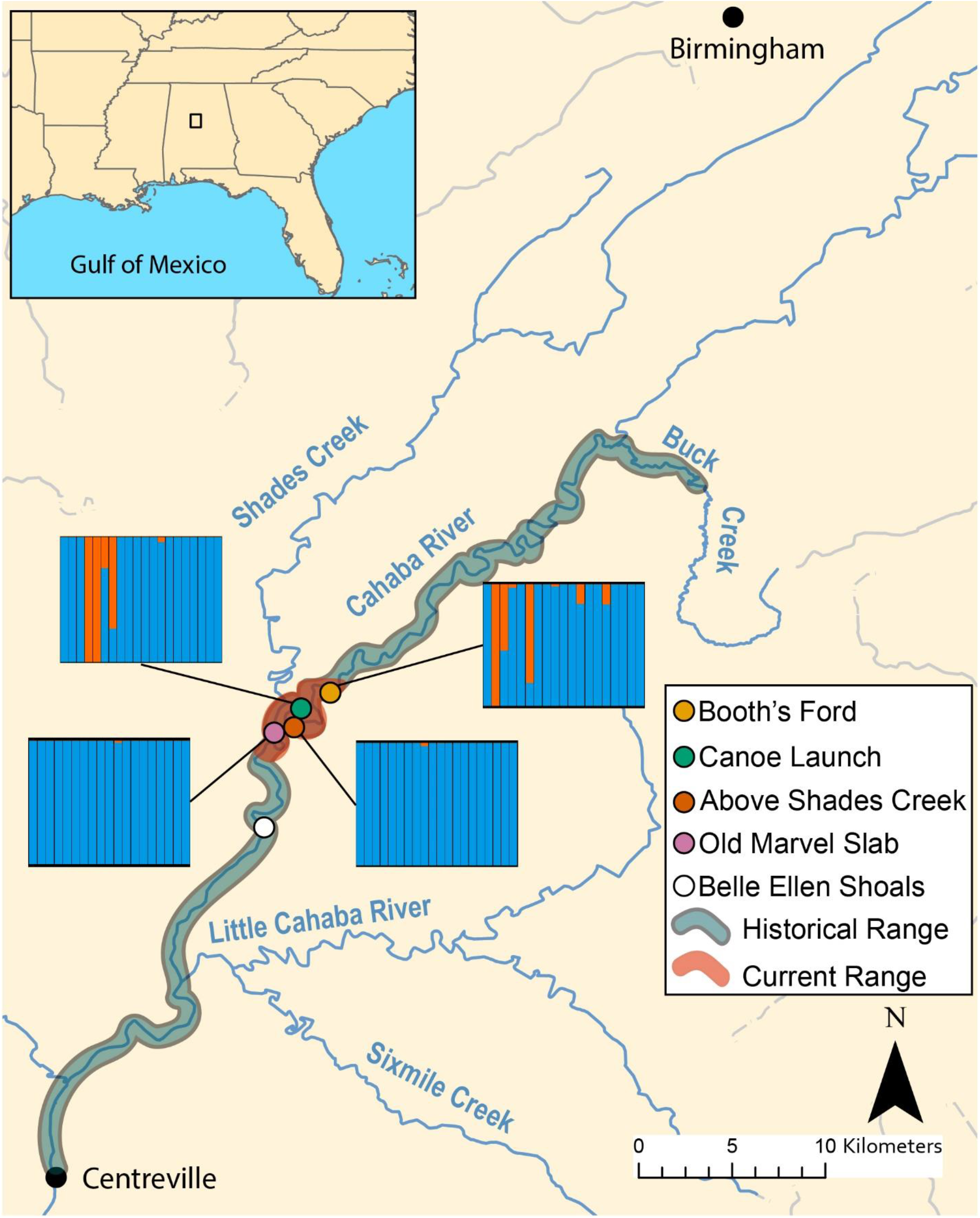
Map of known historical and current range of *L. compacta*, collection sites, and other landmarks. Lines from collection sites lead to ADMXITURE plots with K = 2 for each site. Each column is an individual with ADMIXTURE proportions of the two inferred ancestral populations.

### Molecular data generation

Tissue clips from the foot of 20 individuals per collection site were taken and subjected to a standard proteinase K digestion. DNA was extracted with the Qiagen DNeasy Plant Mini Kit with a minor modification to allow for tissue digested with proteinase K. We used a plant kit because it works well on freshwater gastropods that produce large amounts of mucus polysaccharides (Whelan et al., 2019). The integrity of whole genomic DNA was checked on a 1% agarose gel and quantified with a Qubit DNA assay. Extracted DNA was standardized to 120 ng/μL for 2bRAD library prep.

A reduced representation genomic library was generated for genotyping using the AlfI enzyme and the 2bRAD library prep protocol of Wang et al. (2012). This RAD-seq approach uses a type IIB restriction enzyme that has two recognition sites. AlfI recognizes two sites separated by six base pairs and makes two cuts that each have a one base pair overhang 12 base pairs from the 5’ and 3’ ends of the restriction sites. Following Whelan et al. (2019), we did a 1/16^th^ genomic reduction by using adaptors in the ligation step that had an “NC” overhang, thus only binding to AlfI RAD-loci that had a G base pair at the first base pair of each restriction cut overhang. For more details, see Wang et al. (2012) and the lab protocol on the FigShare repository for this study (DOI: 10.6084/m9.figshare.12014619).

All samples were dual-indexed for pooling. Sequencing occurred in multiple batches. The first batch had 48 *L. compacta* samples pooled in equimolar concentrations and sequenced on a single lane. The other individuals were pooled in equimolar concentrations with samples from projects on conservation genomics of other pleurocerid species, and 87 individuals were sequenced per HiSeq 4000 lane. Although batch effects in RADseq data have been recently noted in studies that used different read lengths among sequencing runs (Leigh, Lischer & Keller, 2018) and in species introgression studies (Lambert et al., 2019), such issues were not relevant to our sequencing design or study objectives. Nevertheless, we took steps to limit potential batch effects by implementing strict filtering parameters during dataset assembly (see below). Pooled libraries were sequenced on an Illumina HiSeq 4000 with 1 X 75bp chemistry at University of Oregon Genomics and Cell Characterization Core Facility.

Raw Illumina reads were demultiplexed with the STACKS 1.48 (Catchen et al., 2013) module *process_radtags*, allowing for one mismatch per barcode. Demultiplexed reads were quality filtered with the script QualFilterFastq.pl (http://github.com/Eli-Meyer/sequence_processing) for any read that had five or more base pairs with Phred quality scores less than 20. Reads were processed with scripts from SHRiMP 2.23 (Rumble et al., 2009) and subsequently trimmed to AlfI RAD-loci with the script AlfIExtract.pl (http://github.com/Eli-Meyer/2bRAD_utilities). As this step removes any sequence that is not part of the RAD-locus, adaptor sequences and non-target sequences are removed from the sequencing reads. RAD-loci, defined as the stretch of DNA cut by the AlfI enzyme, were assembled with the STACKS 1.48 pipeline *denovo_map.pl* as no reference genome is available for *L. compacta*. For *denovo_map.pl* parameters, we set minimum stack depth to five (-m 5), distance allowed between stacks to three (-M 3), and distance between catalog RAD-loci to two (-n 2). These parameters were determined to be most appropriate for our data following Paris et al. (2017). All other *denovo_map.pl* parameters were set to defaults.

After assembly, RAD-loci were filtered for missing data using the STACKS program *populations*. In order to pass filtering steps, a RAD-locus had to be present in 75% of individuals from any given collection site and also present at three collection sites. RAD-loci that had a minimum minor allele frequency of less than 2.5% or heterozygosity higher than 50% were removed to limit the influence of paralogy and misassembly on final datasets. Sequencing coverage of RAD-loci with SNPs was measured with vcftools (Jombart & Ahmed, 2011). Kinship coefficients among individuals were inferred with KING (Manichaikul et al., 2010). Files output by STACKS were formatted for KING with PLINK 1.9 (Chang et al., 2015), and pairwise kinship coefficients were calculated with the KING flag “--kinship”. No individuals were determined to be closely related by KING so no further dataset filtering was done.

After filtering, a dataset that included all SNPs per RAD-locus and a dataset with only one random SNP per RAD-locus were generated. We assume that RAD-loci are unlinked and that the one SNP per RAD-locus dataset had zero linkage disequilibrium. Analyses employed the one SNP per RAD-locus dataset, unless otherwise noted.

### Population genetics analyses

Average observed heterozygosity (H_o_), expected heterozygosity (H_e_), nucleotide diversity (Π), and FIS at each collection site were calculated by *populations*. The number of private alleles at each site was also reported by *populations*. Average allelic richness (A_r_) was calculated with the R (R Core Team, 2020) package diveRsity (Keenan et al., 2013). An analysis of molecular variance (AMOVA; Excoffier, Smouse & Quattro, 1992) was done with the R packages poppr (Kamvar, Tabima & Grünwald, 2014) to test genetic structure among collection sites. AMOVA was implemented with the function “poppr.amova” using the ade4 method (Dray & Dufour, 2007) and 10,000 permutations.

We also tested for a pattern of isolation by distance by measuring the correlation between pairwise F_ST_ values and geographical distance between collection sites. Pairwise F_ST_ values were calculated using the Weir and Cockerham (1984) method with the R package hierfstat (Goudet, 2005). Stream distance was measured in Google Earth by tracing paths between collection sites along the Cahaba River (see Table 2). River distance was used, rather than straight line distances, because migration over land is impossible for gill breathing pleurocerids. A Mantel test was performed with the R package ade4 (Dray & Dufour, 2007), and significance was tested with 1,000 permutations. However, Mantel tests have been criticized as a method for testing isolation by distance (Legendre, Fortin & Borcard, 2015; Meirmans, 2015), so we also performed a multiple regression on distance matrices with 1,000 permutations using the R package ecodist and its MRM function (Goslee & Urban, 2007). In addition to a pattern of isolation by distance, past studies have shown that many freshwater organisms, including pleurocerids, display a pattern of increased genetic diversity in more downstream populations (Paz-Vinas et al., 2015; Whelan et al., 2019). Therefore, to better assess riverscape genetic patterns of *L. compacta*, we performed linear regression of distance from the most downstream site against H_o_, H_e_, A_r_, and Π. Linear regressions were done in R.

**Table 2:** Pairwsie FST and distances (km) between sites. FST below diagonal and distances above diagonal

|  | Booth's Ford | boat launch | above Shades Creek | old Marvel slab |
| --- | --- | --- | --- | --- |
| Cahaba River at Booth's Ford | - | 4.62 | 5.55 | 9.2 |
| Cahaba River at canoe launch | 0.04 | - | 0.98 | 4.57 |
| Cahaba River above Shades Creek | 0 | 0.05 | - | 3.64 |
| Cahaba River at old Marvel slab | 0.03 | 0.04 | 0.03 | - |

We examined clustering of *L. compacta* genetic data with discriminant analysis of principal components (DAPC). We used the multiple SNPs per RAD-locus dataset and the R package adegenet (Jombart & Ahmed, 2011) to perform DAPC. We first used the adegenet function “find.clusters” testing up to 25 clusters and using Bayesian information criteria (BIC) to identify the best-fit number of clusters for our data. Using the number of clusters with the lowest BIC value, we performed a DAPC with the adegenet function “dapc” and plotted the results in R.

We inferred genomic admixture of *L. compacta* individuals with ADMIXTURE 1.3 (Shringarpure et al., 2016). ADMIXTURE assumes zero linkage disequilibrium, so we used the one SNP per RAD-locus dataset. ADMIXTURE analyses were run with the AdmixPipe pipeline (Mussmann et al., 2020). To determine the best-fit number of clusters (K) for our data, K values from 1 to 5 were assessed with 20% cross-validation. Twenty replicates of ADMIXTURE were run at each K, and the best-fit K was determined as the value that had the lowest average CV score across replicates. ADMIXTURE results were visualized with Clumpak (Kopelman et al., 2015).

Genomic co-ancestry among individuals was also assessed with fineRADstructure (Malinsky et al., 2018). Unlike ADMIXTURE, fineRADstructure can use linked SNPs and provides additional information on individual genomic background. Thus, the multiple SNPs per RAD-locus dataset was used for fineRADstructure analyses. First, a co-ancestry matrix was inferred with the script RADpainter. Subsequently, clustering was done with the Markov chain Monte Carlo method of fineRADstructure, running for 500,000 generations and sampling every 1,000 generations; the first 200,000 generations were discarded as burn-in (non-default parameters: -x 200000 -y 300000 -z 1000). We also inferred a tree for visualization with fineRADstructure using the tree-building algorithm of Lawson et al. (2012) with 10,000 attempts (non-default parameters: -m T -x 10000). fineRADstructure results were plotted with R scripts included in the fineRADstructure package.

### Code and data availability

All bash and R scripts used for processing and analyzing data are available at github.com/nathanwhelan. Demultiplexed raw Illumina reads have been uploaded to NCBI under BioProject PRJNA631794. Assembled datasets in various file formats (e.g., vcf, genepop) and the 2bRAD library prep protocol are available on FigShare (DOI: 10.6084/m9.figshare.12014619).

## Results

### Sample Collection

During survey work, we collected *L. compacta* from Cahaba River at old Marvel slab upstream to Cahaba River at Booth’s Ford (Fig. 3). All sites except Cahaba River at Shades Creek are sites where *L. compacta* was not found during survey work over the least 30 years. Our collections represent a 1.83 km downstream range extension and a 4.76 km upstream extension compared to the previously documented contemporary range of *L. compacta* (Whelan, Johnson & Harris, 2012). While this study was ongoing, 3 putative *L. compacta* individuals were collected at Cahaba River at Belle Ellen Shoals (Fig. 3) during a general mollusk survey (Johnson, 2019). However, species identification was uncertain and *L. compacta* appeared exceedingly rare. Therefore, individuals from Cahaba River at Belle Ellen Shoals were not included in our analyses, and we consider this record unconfirmed without additional positive survey results.

### Molecular data and population genetics

DNA yields for two individuals were too low for library preparation so only 19 individuals were sequenced from Cahaba River at old Marvel Slab and Cahaba River at Booth’s Ford. The number of demultiplexed raw reads per individuals varied from 930,062-10,146,649 (mean = 4,836,812). Much of the variation in raw reads can be attributed to whether the individual was sequenced on a HiSeq 4000 lane with 48 or 87 samples. Aside from raw read number, we saw no evidence of batch affects like individuals from one sequencing run all clustering together in analyses (see below). After initial raw-read filtering, the number of reads that passed quality filtering steps ranged from 865,314-9,838,187 (mean = 4,632,510). Assembly with the STACKS *denovo_map* pipeline resulted in 105,542 RAD-loci. Filtering with *populations*, including removal of 4,009 invariant RAD-loci that passed all filters, resulted in a dataset with 4,962 RAD-loci with at least one SNP. Per individual average sequencing coverage of filtered RAD-loci with at least one SNP, excluding missing genotypes, ranged from 31.7-343.2. Average sequencing coverage across variable RAD-loci, excluding missing genotypes, was 163.7. Kinship coefficients inferred with KING indicated that no individuals were closely related (i.e., half or full siblings).

The number of private alleles at each site ranged from 28-262 (Table 1). H_o_ at each collection site ranged from 0.0963-0.1568, and H_e_ ranged from 0.0980 to 0.1801 (Table 1). At each site, H_o_ was lower than H_e_, except at Cahaba River at canoe launch where H_o_ was 0.001 greater than H_e_ (Table 1). The difference between H_o_ and H_e_ was largest at Cahaba River above Shade Creek and Cahaba River at Booth’s Ford. A_r_ and Π ranged from 1.4511-1.8241 and 0.1010-0.1829, respectively (Table 1). F_IS_ values ranged from 0.0134-0.1934 (Table 1), with the highest values being at Cahaba River above Shades Creek and Cahaba River at Booth’s Ford. Overall, genetic diversity was greatest at the most upstream site, Cahaba River at Booth’s Ford, and lowest at the most downstream site, Cahaba River at old Marvel slab. All linear regressions of diversity statistics vs distance from the most downstream site were non-significant (p ≥ 0.169).

Pairwise F_ST_ values among sites ranged from 0.0-0.055 (Table 2). We found no evidence of an isolation by distance pattern among sites (Mantel test, p = 0.843; multiple regression, p = 0.428). According to the AMOVA, significant genetic structure was present among collection sites (p = 0.004), but only 4.16% of variation was explained by collection site. In contrast, 81.8% of genetic variation was explained by within individual variation, further indicating high amounts of gene flow among collection sites.

DAPC indicated two genetic clusters were present in our data. Data were explained by a single discriminate function, and results are therefore presented as a frequency histogram (Fig. 4). ADMIXTURE analyses indicated that genetic diversity from two ancestral populations were present in our data (K = 2). Most individuals from across the range of *L. compacta* had a genomic admixture profile that was dominated by a genomic background from a single ancestral population, likely indicating that overall genomic diversity has been lost across the range of *L. compacta*. Nevertheless, 14 individuals had varying levels of admixture with a second ancestral population (Fig. 3). fineRADstructure analyses corroborated ADMIXTURE analyses as two semi-distinct groupings were recovered by fineRADstructure (Fig. 5, Supplementary Fig. 1). fineRADstructure groupings did not correspond to collection site or any other obvious variable, indicating gene flow among collection sites. Notably, six individuals with comparably high co-ancestry proportions (upper right of co-ancestry matrix in Fig. 5) correspond to individuals in ADMIXTURE analyses with a large proportion of genetic background from the less common ancestral population (represented by orange in Fig. 3).

**Figure 4:**
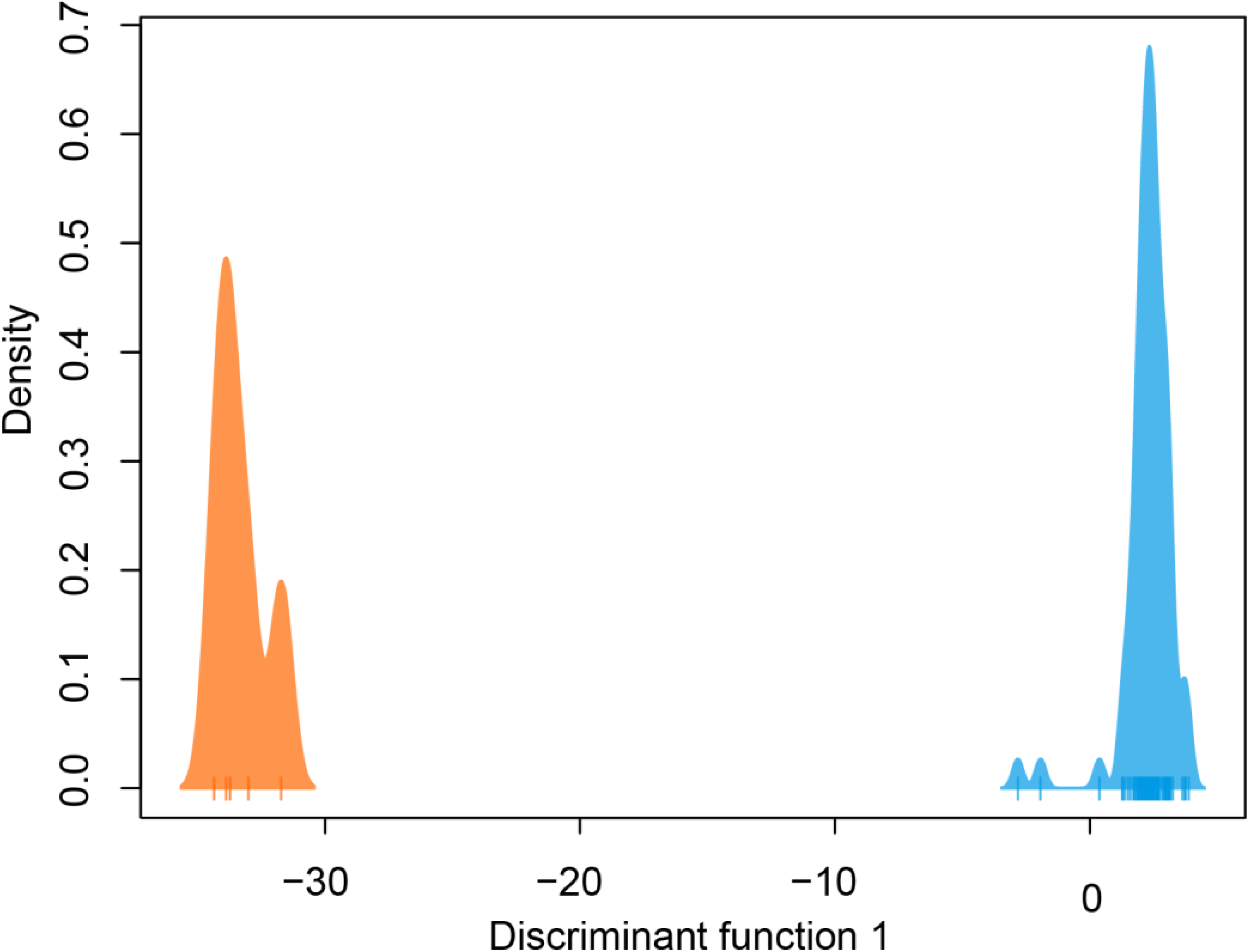
DAPC plot colored by genetic cluster. Tick marks on x-axis represent individuals.

**Figure 5:**
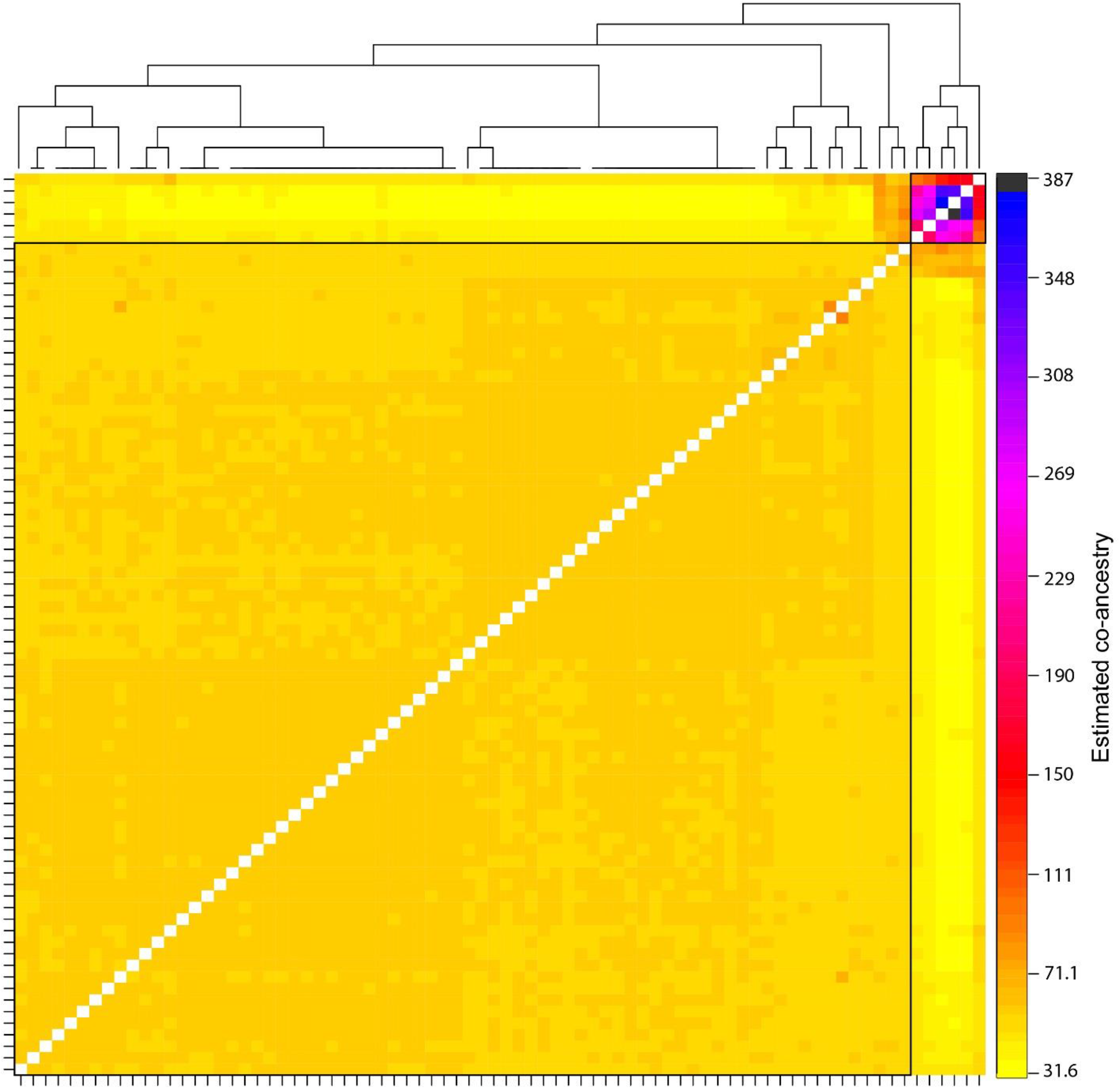
Pairwise co-ancestry matrix and simple tree inferred with fineRADstructure. Boxes surround the two main groupings. Tick marks represent individuals, but labels have been removed for visualization. For a figure with full taxon labels, see Supplementary Figure 1.

## Discussion

Our findings provide reasons to be optimistic about the survival of *L. compacta*. Despite a drastic range reduction in the last 120 years, we found *L. compacta* more widespread than documented in other recent surveys. Furthermore, the remaining sites where *L. compacta* occurs retain a relatively high amount of genetic diversity. Across its range, *L. compacta* had similar levels of H_o_ and Π to *L. ampla*, a species that is currently found across the historical range of *L. compacta* and in some tributaries like Shades Creek and Little Cahaba River (Whelan et al., 2019). The lowest genetic diversity values observed for *L. compacta* were greater than the lowest values determined for *L. ampla*. This observation rejects one of our main hypotheses that *L. compacta* would have lower genetic diversity than the more widespread *L. ampla*. Nevertheless, *L. compacta* is restricted to a 9.2 km stretch of river, and *L. compacta* has likely lost range-wide genetic diversity. This probable loss of evolutionary potential could be detrimental to the long-term survival of the species.

Observed *L. compacta* genetic patterns often conflicted with predictions made by broad-scale hypotheses about riverscape genetics. For example, we did not see an isolation by distance pattern, which is common among freshwater taxa (Hughes, Schmidt & Finn, 2009) and was documented in *L. ampla* (Whelan et al., 2019). We also did not uncover a pattern of increased genetic diversity in downstream populations, despite such a pattern being present in numerous plants and animals (Paz-Vinas et al., 2015), including *L. ampla* (Whelan et al., 2019). Patterns determined for *L. compacta* are likely explained by a drastic range reduction and the limited scale at which we performed the current study. That is, gene flow across the 9.2 km contemporary range of *L. compacta* explains observed patterns of riverscape genetic diversity.

### Genetic diversity across a small landscape

The two most distant collection sites in this study were separated by a smaller distance (9.2 km) than all but two sites sampled for *L. ampla* in a previous study (Whelan et al., 2019). Therefore, it is difficult to make direct comparisons between genetic patterns of *L. ampla* and *L. compacta*. However, we can leverage differences in geographical scale between the two studies to make inferences about fine-scale versus long-distance genetic patterns in pleurocerids. F_ST_ values among *L. compacta* collection sites (Table 2) were much lower than values determined for populations of *L. ampla* (F_ST_ 0.377-0.773; Whelan et al. 2019). Furthermore, even though AMOVA indicated significant genetic structure among *L. compacta* collection sites, the small amount of genetic variation that is explained by collection site probably limits its biological relevance. Overall, these data indicate that pleurocerid riverscape genetic patterns across small distances will not always follow common patterns such as isolation by distance and increased genetic diversity at more downstream collection sites. This is likely attributable to gene flow and random drift that prevent the establishment of genetic patterns typically seen across more geographically separated collection sites. From a historical standpoint, we hypothesize that *L. compacta* previously displayed an isolation by distance pattern across its range, similar to the patterns determined for *L. ampla* (Whelan et al., 2019). We think this scenario is likely given limited dispersal abilities of pleurocerids and patterns established for *L. ampla*, a species that retains a much larger portion of its historic range in the Cahaba River drainage than *L. compacta*. Whether or not there was a similar historical pattern of increased genetic diversity in downstream populations of *L. compacta* is more difficult to infer, as such a pattern may not be influenced solely by dispersal ability.

Given the well-documented decline of *L. compacta*, a small number of individuals with a less common genomic background suggests that the species has lost genetic diversity through bottleneck and drift. Patterns seen in DAPC, ADMIXTURE, and fineRADstructure were not driven by geography as individuals with the less common genomic background were not found in adjacent sites (orange in DAPC and ADMIXTURE plots and upper right corner of fineRADstructure plot; Figs. 3-5, Supplementary Fig. 1). Although individuals with some admixture from the uncommon ancestral population may be present in unsampled individuals at Cahaba River at old Marvel slab and Cahaba River at canoe launch, they would be uncommon. Recent migration is an unlikely explanation of observed co-ancestry profiles as it would indicate that a sizeable population of *L. compacta* exists elsewhere in the Cahaba River. The most likely hypothesis for explaining observed clustering and co-ancestry profiles (Figs. 3-5) is a genetic bottleneck resulting from species decline in the 20^th^ century. In this scenario, *L. compacta* was genetically diverse across its historical range prior to decline, but range contraction caused a considerable loss of genetic diversity. In turn, genetic drift resulted in the observed coancestry pattern of one ancestral population being more common in extant individuals (Figs. 3, 5).

Broadly, genetic structure across the current range of *L. compacta* can be characterized by a single population with some subpopulation structure at Cahaba River above Shades Creek and Cahaba River at Booth’s Ford (Figs. 3-5; Supplementary Figure 1). The subpopulation structure appears to be causing a Wahlund effect (Whalund, 1928). That is, the Wahlund effect predicts the lower H_o_ values compared to H_e_ values and the higher F_IS_ values seen in collection sites with inferred subpopulation structure (Fig. 3; Table 2). An alternative explanation for the observed pattern of F_IS_ and H_e_ is null alleles. However, null alleles are unlikely as they would increase pairwise F_ST_ values (De Meeûs, 2018) that are uniformly low across populations (Table 2). Despite the putative presence of a Wahlund effect, Cahaba River above Shades Creek and Cahaba River at Booth’s Ford have greater genomic diversity than the two other sites (Table 1; Figs. 3-5). These sites may have better habitat suitability than the other two, allowing for *L. compacta* to persist with greater genetic diversity as the species declined in the 20^th^ century.

### *Conservation of* Leptoxis compacta

*Leptoxis compacta* suffered a massive decline during the 20^th^ century, a period of intense mining, forestry, and urban development in the Cahaba River drainage (Onorato, Angus & Marion, 2000; Pitt, 2000; Shepard et al., 1994; Tolley-Jordan, Huryn & Bogan, 2015). The decline was so drastic that *L. compacta* was considered extinct less than a decade ago. Conservation efforts are needed to ensure the long-term survival of *L. compacta* as the species is at risk from both chronic habitat degradation and one-time catastrophic events. Two potential management strategies for *L. compacta* are habitat restoration and reintroduction with captively reared individuals.

In this study, we report an 8.26 km known range extension for *L. compacta*. One site, Cahaba River at old Marvel Slab, was previously the focus of intense habitat restoration through the removal of a low-level dam (Johnson et al. 2013). The site may have also benefited from improved water quality in Shades Creek (ADEM, 2007; ADEM, 2012) as the site is just below its confluence with the Cahaba River. Since removal of the low-level dam, increases in fish abundance and diversity have been reported (Bennett et al., 2015). Considering *L. compacta* was not found at this site by Whelan, Johnson & Harris (2012), we think habitat either improved from a point where *L. compacta* could not survive or from a point of considerably lower carrying capacity. As the only undammed, major river in the southeastern United States, the Cahaba River is much less modified than most other systems in the southeast. Our findings suggest that imperiled gastropods will benefit from water quality and habitat improvements even in relatively “pristine” river systems. Improving habitat, or identifying suitable habitat, will be a necessary starting point for *L. compacta* reintroduction efforts.

In addition to having a small range, *L. compacta* only exists along a single river path. This means that one catastrophic event such as a massive point source pollution event above Cahaba River at Booth’s Ford could result in extinction of *L. compacta*. Such an event is not merely a hypothetical. In 2016, a gasoline pipeline spill came perilously close to the Cahaba River (Pillion, 2016). To mitigate the risks of a single catastrophic event, reintroduction efforts should emphasize range expansion outside the mainstem Cahaba River. Of course, reintroduction efforts also must be limited by the historical range of any given species. Thus, lower Buck Creek is potentially an ideal reintroduction site if habitat quality is sufficient for the persistence of *L. compacta*. Once a suitable reintroduction site is chosen, managers will need to choose a broodstock site. This decision should be informed with genetic data. The absence of an isolation by distance effect across the current range of *L. compacta* indicates that managers do not need to prioritize potential broodstock sites based on whether they are geographically proximate to reintroduction sites. Rather, sites with high genetic diversity and ease of access should be prioritized for broodstock. Therefore, the Cahaba River above Shades Creek is likely an ideal broodstock location. Moreover, *L. compacta* is easy to sample and relatively easy to distinguish from other sympatric species at Cahaba River above Shades Creek, making it ideal from both a genetic and sampling standpoint.

## Conclusions

Even though *L. compacta* was considered extinct less than a decade ago, we now know more about this species than most other freshwater gastropods. This is helpful for conservation of *L. compacta* as the biggest barrier to effective management strategies for most freshwater gastropods is a lack of data. Future research efforts should focus on differences in dispersal dynamics among pleurocerids and causes of differences in riverscape genetic patterns seen between *L. ampla* and *L. compacta*. As more population genomic data becomes available for pleurocerids, we will be better suited to develop strategies to conserve these critically important components of many North American riverine ecosystems.

## Acknowledgements

We thank Lauren Allred, Jecca Shumante, and Living River: A Retreat on the Cahaba for providing access to field sites. Haley Dutton (Auburn University) assisted in the field. Jeffrey Garner (Alabama Department of Conservation and Natural Resources), Matthew Galaska (University of Washington), Kenneth Halanych (Auburn University) and Katherine Bockrath (U.S Fish and Wildlife Service) provided advice on multiple components of this work. Rodolfo Jaffé, Brian Hand, and an anonymous reviewer provided comments that improved the manuscript. The findings and conclusions in this article are those of the authors and do not necessarily represent the views of the U.S. Fish and Wildlife Service.

---

**Supplementary Table 1:** Collection localities, Auburn University Museum of Natural History, and SRA accession numbers.

| Lab Number | Locality | AUMNH Catalog Number | NCBI SRA accession |
| --- | --- | --- | --- |
| Lcom_pop01 | Cahaba River at old Marvel slab | 45652 | SRR11773571 |
| Lcom_pop02 | Cahaba River at old Marvel slab | 45653 | SRR11773570 |
| Lcom_pop03 | Cahaba River at old Marvel slab | 45654 | SRR11773559 |
| Lcom_pop04 | Cahaba River at old Marvel slab | 45655 | SRR11773548 |
| Lcom_pop05 | Cahaba River at old Marvel slab | 45656 | SRR11773537 |
| Lcom_pop06 | Cahaba River at old Marvel slab | 45657 | SRR11773604 |
| Lcom_pop08 | Cahaba River at old Marvel slab | 45658 | SRR11773593 |
| Lcom_pop09 | Cahaba River at old Marvel slab | 45659 | SRR11773582 |
| Lcom_pop10 | Cahaba River at old Marvel slab | 45660 | SRR11773573 |
| Lcom_pop11 | Cahaba River at old Marvel slab | 45661 | SRR11773572 |
| Lcom_pop12 | Cahaba River at old Marvel slab | 45662 | SRR11773569 |
| Lcom_pop13 | Cahaba River at old Marvel slab | 45663 | SRR11773568 |
| Lcom_pop14 | Cahaba River at old Marvel slab | 45664 | SRR11773567 |
| Lcom_pop15 | Cahaba River at old Marvel slab | 45665 | SRR11773566 |
| Lcom_pop16 | Cahaba River at old Marvel slab | 45666 | SRR11773565 |
| Lcom_pop17 | Cahaba River at old Marvel slab | 45667 | SRR11773564 |
| Lcom_pop18 | Cahaba River at old Marvel slab | 45668 | SRR11773563 |
| Lcom_pop19 | Cahaba River at old Marvel slab | 45669 | SRR11773562 |
| Lcom_pop20 | Cahaba River at old Marvel slab | 45670 | SRR11773561 |
| Lcom_pop21 | Cahaba River at above Shades Creek | 45671 | SRR11773560 |
| Lcom_pop22 | Cahaba River at above Shades Creek | 45672 | SRR11773558 |
| Lcom_pop23 | Cahaba River at above Shades Creek | 45673 | SRR11773557 |
| Lcom_pop24 | Cahaba River at above Shades Creek | 45674 | SRR11773556 |
| Lcom_pop25 | Cahaba River at above Shades Creek | 45675 | SRR11773555 |
| Lcom_pop26 | Cahaba River at above Shades Creek | 45676 | SRR11773554 |
| Lcom_pop27 | Cahaba River at above Shades Creek | 45677 | SRR11773553 |
| Lcom_pop28 | Cahaba River at above Shades Creek | 45678 | SRR11773552 |
| Lcom_pop29 | Cahaba River at above Shades Creek | 45679 | SRR11773551 |
| Lcom_pop30 | Cahaba River at above Shades Creek | 45680 | SRR11773550 |
| Lcom_pop31 | Cahaba River at above Shades Creek | 45681 | SRR11773549 |
| Lcom_pop32 | Cahaba River at above Shades Creek | 45682 | SRR11773547 |
| Lcom_pop33 | Cahaba River at above Shades Creek | 45683 | SRR11773546 |
| Lcom_pop34 | Cahaba River at above Shades Creek | 45684 | SRR11773545 |
| Lcom_pop35 | Cahaba River at above Shades Creek | 45685 | SRR11773544 |
| Lcom_pop36 | Cahaba River at above Shades Creek | 45686 | SRR11773543 |
| Lcom_pop37 | Cahaba River at above Shades Creek | 45687 | SRR11773542 |
| Lcom_pop38 | Cahaba River at above Shades Creek | 45688 | SRR11773541 |
| Lcom_pop39 | Cahaba River at above Shades Creek | 45689 | SRR11773540 |
| Lcom_pop40 | Cahaba River at above Shades Creek | 45690 | SRR11773539 |
| Lcom_pop41 | Cahaba River at Booth's Ford | 45691 | SRR11773538 |
| Lcom_pop42 | Cahaba River at Booth's Ford | 45692 | SRR11773536 |
| Lcom_pop44 | Cahaba River at Booth's Ford | 45693 | SRR11773613 |
| Lcom_pop45 | Cahaba River at Booth's Ford | 45694 | SRR11773612 |
| Lcom_pop46 | Cahaba River at Booth's Ford | 45695 | SRR11773611 |
| Lcom_pop47 | Cahaba River at Booth's Ford | 45696 | SRR11773610 |
| Lcom_pop48 | Cahaba River at Booth's Ford | 45697 | SRR11773609 |
| Lcom_pop49 | Cahaba River at Booth's Ford | 45698 | SRR11773608 |
| Lcom_pop50 | Cahaba River at Booth's Ford | 45699 | SRR11773607 |
| Lcom_pop51 | Cahaba River at Booth's Ford | 45700 | SRR11773606 |
| Lcom_pop52 | Cahaba River at Booth's Ford | 45701 | SRR11773605 |
| Lcom_pop53 | Cahaba River at Booth's Ford | 45702 | SRR11773603 |
| Lcom_pop54 | Cahaba River at Booth's Ford | 45703 | SRR11773602 |
| Lcom_pop55 | Cahaba River at Booth's Ford | 45704 | SRR11773601 |
| Lcom_pop56 | Cahaba River at Booth's Ford | 45705 | SRR11773600 |
| Lcom_pop57 | Cahaba River at Booth's Ford | 45706 | SRR11773599 |
| Lcom_pop58 | Cahaba River at Booth's Ford | 45707 | SRR11773598 |
| Lcom_pop59 | Cahaba River at Booth's Ford | 45708 | SRR11773597 |
| Lcom_pop60 | Cahaba River at Booth's Ford | 45709 | SRR11773596 |
| Lcom_pop61 | Cahaba River at Lebron canoe launch | 45710 | SRR11773595 |
| Lcom_pop62 | Cahaba River at Lebron canoe launch | 45711 | SRR11773594 |
| Lcom_pop63 | Cahaba River at Lebron canoe launch | 45712 | SRR11773592 |
| Lcom_pop64 | Cahaba River at Lebron canoe launch | 45713 | SRR11773591 |
| Lcom_pop65 | Cahaba River at Lebron canoe launch | 45714 | SRR11773590 |
| Lcom_pop66 | Cahaba River at Lebron canoe launch | 45715 | SRR11773589 |
| Lcom_pop67 | Cahaba River at Lebron canoe launch | 45716 | SRR11773588 |
| Lcom_pop68 | Cahaba River at Lebron canoe launch | 45717 | SRR11773587 |
| Lcom_pop69 | Cahaba River at Lebron canoe launch | 45718 | SRR11773586 |
| Lcom_pop70 | Cahaba River at Lebron canoe launch | 45719 | SRR11773585 |
| Lcom_pop71 | Cahaba River at Lebron canoe launch | 45720 | SRR11773584 |
| Lcom_pop72 | Cahaba River at Lebron canoe launch | 45721 | SRR11773583 |
| Lcom_pop73 | Cahaba River at Lebron canoe launch | 45722 | SRR11773581 |
| Lcom_pop74 | Cahaba River at Lebron canoe launch | 45723 | SRR11773580 |
| Lcom_pop75 | Cahaba River at Lebron canoe launch | 45724 | SRR11773579 |
| Lcom_pop76 | Cahaba River at Lebron canoe launch | 45725 | SRR11773578 |
| Lcom_pop77 | Cahaba River at Lebron canoe launch | 45726 | SRR11773577 |
| Lcom_pop78 | Cahaba River at Lebron canoe launch | 45727 | SRR11773576 |
| Lcom_pop79 | Cahaba River at Lebron canoe launch | 45728 | SRR11773575 |
| Lcom_pop80 | Cahaba River at Lebron canoe launch | 45729 | SRR11773574 |

**Supplementary Figure 1:**
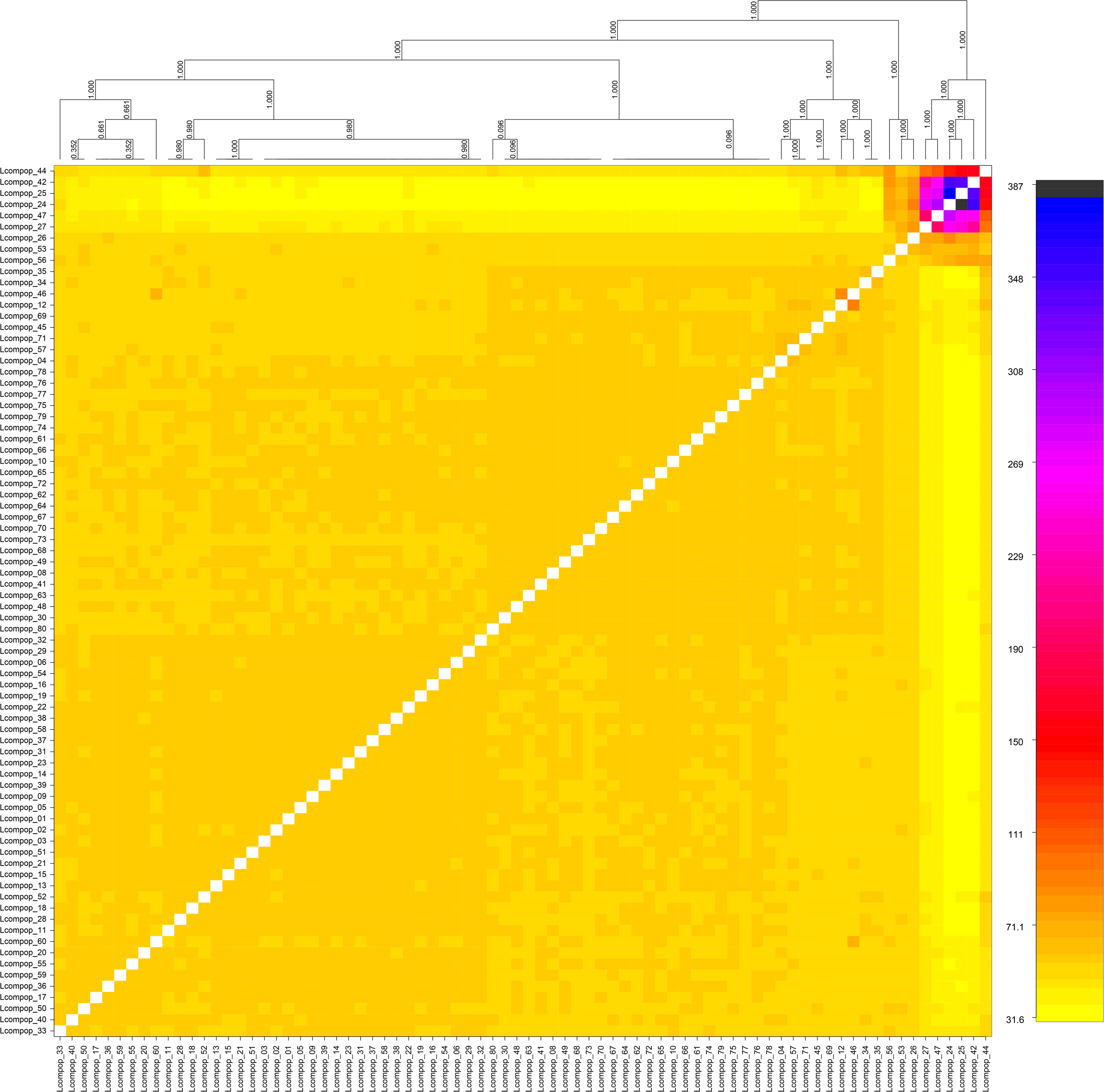
Pairwise co-ancestry matrix and simple tree inferred with fineRADstructure. Tick marks represent individuals. Lcompop_01-20: Cahaba River at old Marvel slab; Lcompop_21-40: Cahaba River above Shades Creek; Lcompop_41-60: Cahaba River at Booth’s Ford; Lcompop_61-80: Cahaba River at canoe launch

